# THE STABILIZATAION OF C-MYC BY THE NOVEL CELL CYCLE REGULATOR, SPY1, DECREASES EFFICACY OF BREAST CANCER TREATMENTS

**DOI:** 10.1101/2022.03.11.483990

**Authors:** Rosa-Maria Ferraiuolo, Bre-Anne Fifield, Caroline Hamm, Lisa Ann Porter

## Abstract

**Purpose:** c-Myc is frequently upregulated in breast cancers, however, targeting c-Myc has proven to be a challenge. Targeting of downstream mediators of c-Myc, such as the ‘cyclin-like’ cell cycle regulator Spy1, may be a viable therapeutic option in a subset of breast cancer subtypes.

**Methods:** Mouse mammary tumour cells isolated from MMTV-Myc mice and human breast cancer cell lines were used to manipulate Spy1 levels followed by tamoxifen or chemotherapeutic treatment with a variety of endpoints. Patient samples from TNBC patients were obtained and constructed into a TMA and stained for c-Myc and Spy1 protein levels.

**Results:** Over time, MMTV-Myc cells show a decreased response to tamoxifen treatment with increasing levels of Spy1 in the tamoxifen-resistant cells. shRNA against Spy1 re-establishes tamoxifen sensitivity. Spy1 was found to be highly elevated in human TNBC cell and patient samples, correlating to c-Myc protein levels. c-Myc was found to be stabilized by Spy1 and knocking down Spy1 in TNBC cells shows a significant increase in response to chemotherapy treatments.

**Conclusions:** Understanding the interplay between protein expression level and response to treatment is a critical factor in developing novel treatment options for breast cancer patients. These data have shown a connection between Spy1 and c-Myc protein levels in more aggressive breast cancer cells and patient samples. Furthermore, targeting c-Myc has proven difficult, these data suggest targeting Spy1 even when c-Myc is elevated can confer an advantage to current chemotherapies.

## INTRODUCTION

Breast cancer is a diverse set of diseases classified largely by the presence or absence of hormone/growth factor receptors. Triple negative breast cancer (TNBC) is identified by the lack of expression of estrogen receptor alpha (ERα), progesterone receptor (PR), and the Her2/neu protein and fall under 5 different genomic subgroups [1-5]. Patients who lack ERα are not candidates for hormone therapy and generally have a less favourable prognosis [6]. A 24.6% change in ERα expression from positive to negative, after relapse was detected in 459 patients [7]. While the mechanism for this downregulation remains to be fully elucidated, ERα levels can be manipulated epigenetically with histone modification and DNA methylation [8]. Understanding the molecular pathways regulating the expression of ERα may provide novel mechanisms of sensitising ERα-negative patients, including those that are triple negative, to available therapies.

Gene amplification and overexpression of the transcription factor c-Myc has been shown in an array of aggressive breast cancers of varying receptor/growth factor status [2, 9-11]. Several studies show that c-Myc overexpression occurs frequently in ERα-negative breast cancers [12, 13] and is implicated as an important driver of TNBC [1, 14, 15]. Approximately 20% of c-Myc amplification in breast cancer is detectable at the mRNA level, implicating regulation at the RNA stability, translation or protein level [16]. In the breast, c-Myc has multiple functions in normal development and is implicated in the initiation and progression of breast cancer [17-20]. Long-term overexpression of c-Myc has become a suggested mechanism towards the development of estrogen-independence [13]. Increased transcriptional activity of c-Myc mimics ERα response to estradiol, specifically through the activation of ERα-targeted genes [21, 22]. This study investigates the implications of persistent c-Myc signalling on ERα status.

While c-Myc is an attractive target for several forms of cancer, mechanistically achieving this goal has been a challenge due to the potent effects of c-Myc on cell proliferation, differentiation, apoptosis, and senescence [23-26]. The dominant negative mutant, Omomyc, has the potential to regress *in vivo* tumours, but tumours quickly re-establish when Omomyc is removed [27]. Targeting c-Myc effectors has demonstrated some effectiveness, of particular interest, inhibiting the G2/M cyclin dependent kinase CDK1 in TNBC has shown impressive synthetic lethality in cell systems [15, 28]. CDK1 is critical for cell proliferation limiting the ability to achieve a safe therapeutic window; hence, exploiting ways to target specific aspects of CDK1 driven activity is required to make this a viable strategy.

One aspect of CDK inhibitors that has been overlooked in the clinic is the presence of ‘cyclin-like’ proteins capable of binding and activating CDKs in the presence of traditional inhibitors [29]. Speedy (Spy1; gene *SPDYA*) is one member of this family. Spy1 can bind and activate G1/S and G2/M CDKs independent of canonical post-translational modifications to enhance proliferation and promote the degradation of p27^Kip1^ [30, 31]. Spy1-bound CDKs have a different conformation with unique substrate preferences [30, 32]. Spy1 levels are low in most adult tissues, selectively expressed in regenerative populations and elevated downstream of c-Myc in several human cancers, including invasive ductal carcinoma of the breast [33-40]. Hence, Spy1-CDKs may represent a unique mechanism to target for selective treatment of specific cancers. Spy1 levels specifically override apoptosis following DNA damage and this mechanism may be particularly potent in drug resistant tumours [41, 42].

Herein, we demonstrate that persistent elevation of c-Myc correlates with increased expression of Spy1 and a reduction in ERα levels. Knockdown of Spy1 reduces c-Myc levels and enhances sensitivity to both hormone and chemotherapy treatment. Hence, specifically targeting Spy1-directed CDKs may be an effective strategy in ERα-negative breast cancers with elevated c-Myc.

## MATERIALS AND METHODS

### Cell Culture

Primary MMTV-Myc tumour cells were isolated from tumours of MMTV-Myc mice and were subcultured in DMEM-F12 media supplemented with 10% FBS, 30,000 units penicillin/streptomycin solution, 5ng/ml EGF, 0.5 µg/ml Hydrocortisone, and 5 µg Insulin. HEK-293, MCF7, MDA-MB-231, and MDA-MB-468 cells were purchased from ATCC and were subcultured in DMEM media containing 10% FBS and 30,000 units penicillin streptomycin solution. LCC9 cells (Lombardi Comprehensive Cancer Center, Georgetown University) were routinely subcultured in DMEM phenol red free media supplemented with 1 mM L-glutamine, 30,000 units penicillin/streptomycin, and 10% charcoal treated FBS. All cells were maintained under normoxic conditions (5% CO_2_) at 37°C.

### Plasmids

Creation of the Myc-Spy1-pCS3 vector was described previously [40]. Rc-CMV-Cyclin E (#8963) and lentiviral constructs pLB (#11619) and pLKO (#8453) were purchased from Addgene. shRNA oligos for Spy1 and a scrambled control were ligated into the pLKO and pLB vectors, as previously described [37].

### Immunoblotting (IB)

Whole cell lysates were prepared as described previously [33] and lysates containing 80-100 µg of protein were subjected to electrophoresis on denaturing 10% SDS polyacrylamide gels and transferred to PVDF-Plus 0.45 micron transfer membrane (Osmonics Inc.) for 2 hours at 30 volts via wet transfer method. Blots were blocked for 1 hour in 1% BSA solution at room temperature. Primary antibodies were reconstituted in blocker: actin MAB150 1R (Chemicon-Millipore; 1:1000), Spy1 (ThermoScientific; 1:1000), c-Myc (Sigma; 1:1000), anti-phospho-c-Myc-S62 (Abcam; 1:1000), anti-phospho-c-Myc-T58 (Santa Cruz Biotechnology; 1:1000), Cyclin E1 (Abcam; 1:1000), anti-p-ERK 1/2 [Thr 202/Tyr 204] (Cell Signaling; 1:1000), and anti-ERK1/2 (Santa Cruz Biotechnology; 1:1000). Secondary antibodies were used at 1:10000 dilution in blocker for 1 hour at room temperature. Chemiluminescence was quantified on an AlphaInnotech HD2 (Fisher) using AlphaEase FC software.

### Transfection/Lentiviral Infection

#### Transfection

Cells were transfected using PEI branched reagent. In brief, 10 µg of DNA was mixed with 30 µg of PEI for 10 minutes then added to a 10 cm tissue culture plate. Transfection media was changed after 24 hours. *Lentiviral Infection:* 8000 cells were seeded in fully supplemented growth media in 96-well plates for 2 hours. Cells were starved by removing serum and penicillin/streptomycin from the media, followed by the use of 1 mg/ml polybrene (Santa Cruz Biotechnology) and MOI 3 of the specific vector used. Infected media was changed to fully supplemented media 24 hours after infection.

### Drug Treatments

Treatments included vehicle control dimethyl sulfoxide (DMSO), 100 nM 4-Hydroxytamoxifen (Sigma), 25 nM doxorubicin (Sigma), 100nM paclitaxel, 6mM cyclophosphamide (Sigma), 25 µM Nu-2058 (Santa Cruz), or 20 µM Roscovitine (Santa Cruz) for specified time points.

### Proliferation Assay

Cells were seeded at 5×10^4^ cell density in 24-well plate. Following treatments cells were collected at specified time points, pelleted and resuspended in 1mL of media. 10 µl sample was collected, trypan blue was added and counted using a haemocytometer.

### BrdU Assay

BrdU stock (10 mM) was dissolved in 10 mL of culture medium to produce a 10 µM labelling solution. Infected and non-infected cells were seeded at 8×10^3^ cell density in a 96-well plate, after 24 hours the culture medium was replaced with BrdU labelling solution for 30 minutes in CO_2_ incubator at 37°C. Cells were washed 2 times with 1x PBS for 2 minutes each. Cells were fixed in 3.7% formaldehyde in PBS for 30 minutes at room temperature. Cells were washed with 1xPBS 3 times for 2 minutes each. Cells were immersed in 0.07N NaOH for 2 minutes, then in 1xPBS (pH 8.5). 20 µl anti-BrdU (Becton Dickinson) was mixed with 50 µl of 0.1% Tween 20/PBS. Cells were incubated for 30 minutes in a humidified chamber with the diluted unconjugated anti-BrdU. Cells were washed with 1xPBS. 50 µl 0.1% Tween 20/PBS was added to cells. Alexa mouse (1:1000) was added for 30 minutes at room temperature. Cells were washed with 1xPBS and incubated with Hoechst. Cells were washed with water and air dried.

### Cyclohexamide (CHX) Treatment

After transfection cells were treated with 50 µg/mL cycloheximide (Sigma) to block *de novo* protein synthesis. After 0.25 to 2 hours, cells were pelleted, harvested and subjected to SDS-PAGE analysis and IB.

### Cell Sorting

Cells were collected, and surface and intracellular staining was performed as described in BD Pharmigen protocol. Briefly, surface stain was performed by the addition of mAB ERα Alexa Fluor 488 antibody (ab194150) for 30 minutes at 4°C, followed by washes with cold stain buffer. Solution was centrifuged, and pellet was resuspended and incubated with Fix/Perm Solution for 50 minutes, followed by washes and intracellular stain with mAB ERα Alexa Fluor 488 antibody (ab194150). Incubation for intracellular stain is 50 minutes at 4°C. Stain was washed prior to running the samples on a BD FACSAria Fusion at excitation 495 nm, emission 519 nm.

### Tissue Microarray (TMA) Construction

Embedded tissue samples were received from Windsor Regional Cancer Centre and constructed into TMAs using an Arraymold Inc., TMA construction and sectioning was completed was described [43].

### Immunohistochemistry (IHC)

TMA sectioned slides were deparaffinized and rehydrated using xylene and decreasing concentrations of EtOH. Slides were washed in 1x PBS. Sodium citrate (pH 6.0) antigen retrieval was performed. Slides were washed in distilled water in 1x PBS. Slides were blocked for endogenous peroxidases using 90 ml methanol/10 ml 30% H_2_O_2_ at room temperature, followed by washes in 1x PBS. Blocking buffer (3%BSA/0.1% Tween in 1xPBS) was added for 1 hour at room temperature in a humidified chamber. Primary antibody (1:200) was added to slides overnight at 4°C. Biotinylated secondary antibody (1:500) was added to the slides for 1 hour at room temperature in a humidified chamber. Vectastain ABC Reagent and DAB Peroxidase Substrate (Vector Labs) were used as per manufacturer’s instructions. Haematoxylin was used as a counterstain. Slides were dehydrated in increasing EtOH concentrations and immersed in xylene.

### Microscopy

Slides were imaged using Leica Stereoscope M205FA. Using Leica LAS V4.3 program, images were taken at 79.7x magnification using 1x stereoscope objective and 159x magnification using 2x stereoscope objective.

### TMA Quantification

Quantification of Spy1 and c-Myc immunostaining intensity was performed using Adobe Photoshop CC 2014 (Adobe Systems Inc. San Jose, CA) using nine random samplings of 10 × 10 pixels each, based on a previously reported densitometry method [44, 45].

## RESULTS

### Primary MMTV-Myc cells acquire resistance to tamoxifen over time in culture

Primary MMTV-Myc cells were passaged over time and their response to tamoxifen over 24 hours measured using BrdU analysis. In early passages (P10-P35) tamoxifen treatment reduced the number of cells going through DNA synthesis by 80-90%, but in late passages (P45-P85) tamoxifen had no significant effect (Figure 1A). Early passage cells (P34) decrease in cell number by ∼35% in response to treatment, while late passage cells (P80) do not respond to tamoxifen treatment (Figure 1B). We then used this model to determine any effects on ERα levels with persistent c-Myc signalling (Figure 1C). Protein levels of ERα begin to decrease dramatically between P30-P80. As shown previously, levels of Spy1 are high in the presence of c-Myc signalling [38], and we show that they remain at an elevated level at all passages (Figure 1C). We sorted primary myc cells for the presence or absence of ER (Figure 1D). All passages had a significantly higher percentage of cells with ER than those that were negative for ER (Figure 1D). Sorted cells were then tested for protein levels. We found that Spy1 levels were significantly elevated in P34 and P80 ER negative populations, while c-Myc was only significantly elevated in P80 ER negative cell populations (Figure 1E). pERK:Total ERK levels remained constant throughout all cell populations at all passages (Figure 1E).

**Figure 1.**
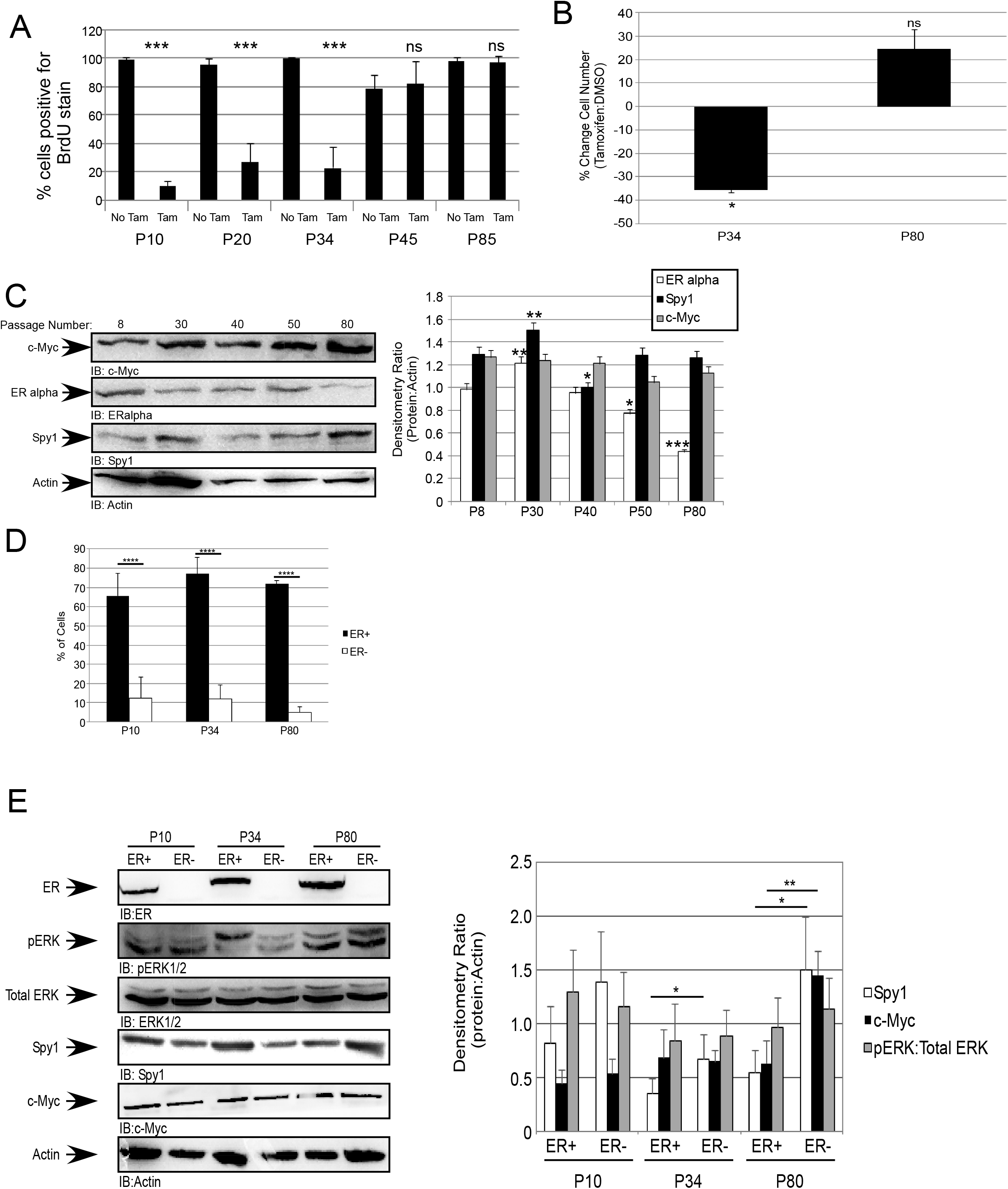
Primary MMTV-Myc cells acquire resistance to tamoxifen over time in culture. (A) Cells at passage (P) 10 to 80 were treated with 100 nM of tamoxifen or vehicle control (DMSO); followed by BrdU analysis. The percent BrdU positive cells are depicted on y-axis. (B) Cells at various passages (denoted on x-axis) were treated with 100 nM of tamoxifen or vehicle control (DMSO) followed by trypan blue exclusion assay. Percent difference between tamoxifen treated and DMSO control depicted on y-axis. (C) SDS-PAGE and IB performed on cells (passage number denoted on the top of representative blot-left panel, and on x-axis of densitometry graph-right panel). (D) Cells at P10, P34 and P80 were sorted on the basis of ERα expression. Quantification of the percentage of ERα shown. (E) SDS-PAGE and IB performed on cells sorted on basis of ERα expression. Representative blot depicted in left panel and quantification of densitometry depicted on right panel. (A-E) Error bars reflect SE between triplicate experiments. Student’s t-test was performed; *p<0.05, **p<0.01, ***p<0.001.

### Spy1 levels affect cellular response to tamoxifen treatment

To determine if Spy1 levels could affect the treatment response of late passage cells (P80), which showed an acquired resistance to tamoxifen, we infected P80 cells with either shScrambled (pLB) or Spy1 knockdown (shSpy1). Knockdown of Spy1 significantly decreased c-Myc protein levels (Figure 2A). Cells with Spy1 knockdown had a significant reduction in cell proliferation over time compared to the control population (Figure 2B). Spy1 knockdown and control cells were treated with 100 nM tamoxifen and viability measured. After 24 hours, Spy1 knockdown significantly decreased the number of viable cells by ∼33% in response to tamoxifen (Figure 2C). This was also reflected by a significant difference in BrdU incorporation in P80-Spy1 knockdown vs. control knockdown cells treated with tamoxifen (Figure 2D).

**Figure 2.**
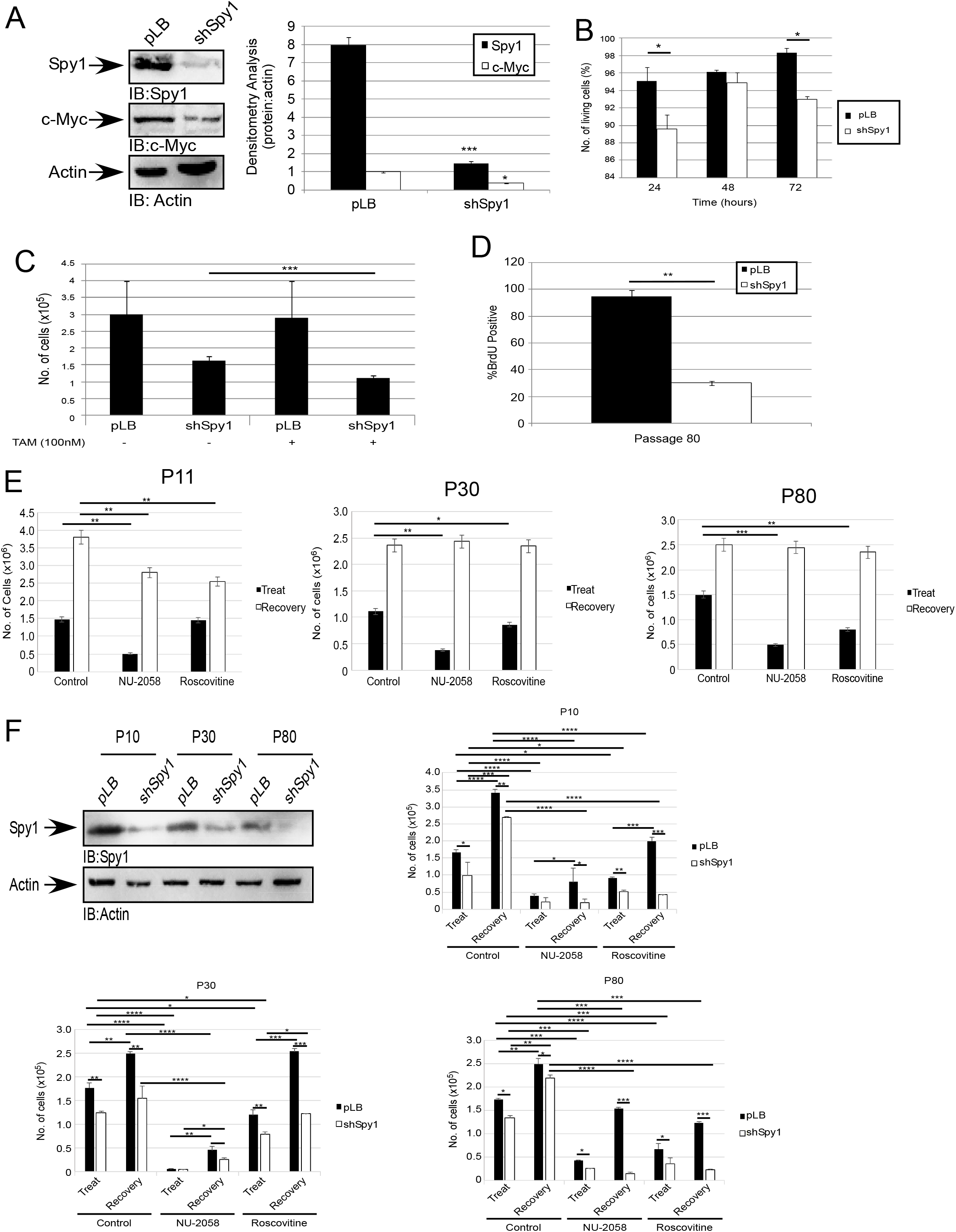
Spy1 levels affect cellular response to tamoxifen. Cells were infected with pLB empty vector or pLB-shSpy1 vector. (A) SDS-PAGE and IB was performed. A representative blot is shown (left panel) and densitometry ratio of protein to loading control actin is shown (right panel). (B) Infected cells were subjected to trypan blue exclusion assay for the indicated time course. (C) Infected cells were treated with 100 nM of tamoxifen or vehicle control (DMSO). Proliferation after treatment was assayed using trypan blue exclusion assay. (D) BrdU analysis of P80 cells infected with pLB or shSpy1. The percent of BrdU positive cells is depicted on the y-axis. (E) Cells at P11 (left panel), P30 (middle panel) and P80 (right panel) were treated with 25µM NU-2058, 20µM roscovitine or vehicle control (DMSO). Proliferation after treatment and after 48 hours recovery following treatment was assessed using trypan blue exclusion assay. (F) Infected cells were treated with 25µM NU-2058, 20µM roscovitine or vehicle control (DMSO) and proliferation assessed via trypan blue exclusion assay following treatment and a 48 hour recovery period. (A-F) Error bars reflect SE between triplicate experiments. Student’s t-test was performed; *p<0.05, **p<0.01,***p<0.001.

Since most c-Myc driven tumours fall in the subcategory of TNBC, and synthetic CDK inhibitors have started to be used as a treatment option to induce synthetic lethality, we tested whether primary myc cells at early to late passages could convey synthetic lethality. Two CDK inhibitors were used; Roscovitine, a pan-CDK inhibitor, and NU-2058, a CDK2 selective inhibitor. All passages responded significantly to both CDK inhibitors, responding best to NU-2058 (Figure 2E). To determine if there were lasting effects after CDK inhibitor treatment, media was changed to contain no inhibitor and the cells were left to recover. Cells were counted 48 hours after media release. All cells recovered; however, only P11 cell numbers remained significantly lower than control while P30 and P80 cell numbers grew faster after the removal of treatment (Figure 2E). We next investigated whether Spy1 levels affect this observation by knocking down Spy1 in each passage followed by treatment with each inhibitor. Spy1 knockdown alone significantly decreased total cell number and the addition of inhibitor treatment further decreased the total cell number, with a greater decrease seen with the selective CDK2 inhibitor (Figure 2F). Furthermore, recovery after treatment showed an increase in cell number in all pLB control cells with and without treatment, whereas a less significant increase in cell number was seen with Spy1 knock down (Figure 2F). This data indicates Spy1 levels may contribute to resistant Myc-driven tumours.

### Spy1 has a role in stabilizing c-Myc

TNBC is associated with elevated levels of c-Myc [28]. We tested whether Spy1 levels correlated with that of c-Myc in TNBC cells (MDA-MB-231 and MDA-MB-468) as compared to ERα-positive MCF7 or their hormone resistant counterpart LCC9 (Figure 3A) [46]. TNBC cells have higher levels of both Spy1 and c-Myc as compared to ERα positive breast cancer cell lines. Interestingly, LCC9 resistant cells also have higher levels of both c-Myc and Spy1 than hormone sensitive MCF7 cells. To elucidate whether Spy1 is essential for elevated c-Myc levels, we infected MDA-MB-231 cells with shScrambled or shSpy1 and collected 24 to 72 hours after infection. Spy1 knockdown significantly reduced c-Myc protein levels (Figure 3B). To determine if Spy1 overexpression can affect the stabilization of c-Myc protein, HEK-293 cells were manipulated to overexpress Spy1 or an empty vector control (pCS3) followed by treatment cycloheximide to block *de novo* protein synthesis and c-Myc protein half-life (t_1/2_) was monitored by immunoblotting over time. Spy1 increases the half-life of c-Myc almost 2-fold (Figure 3C). Half-life of the phosphorylated form was shorter than that of overall levels of c-Myc but also demonstrates a significant increase in the presence of Spy1 (Figure 3C, lower graph).

**Figure 3.**
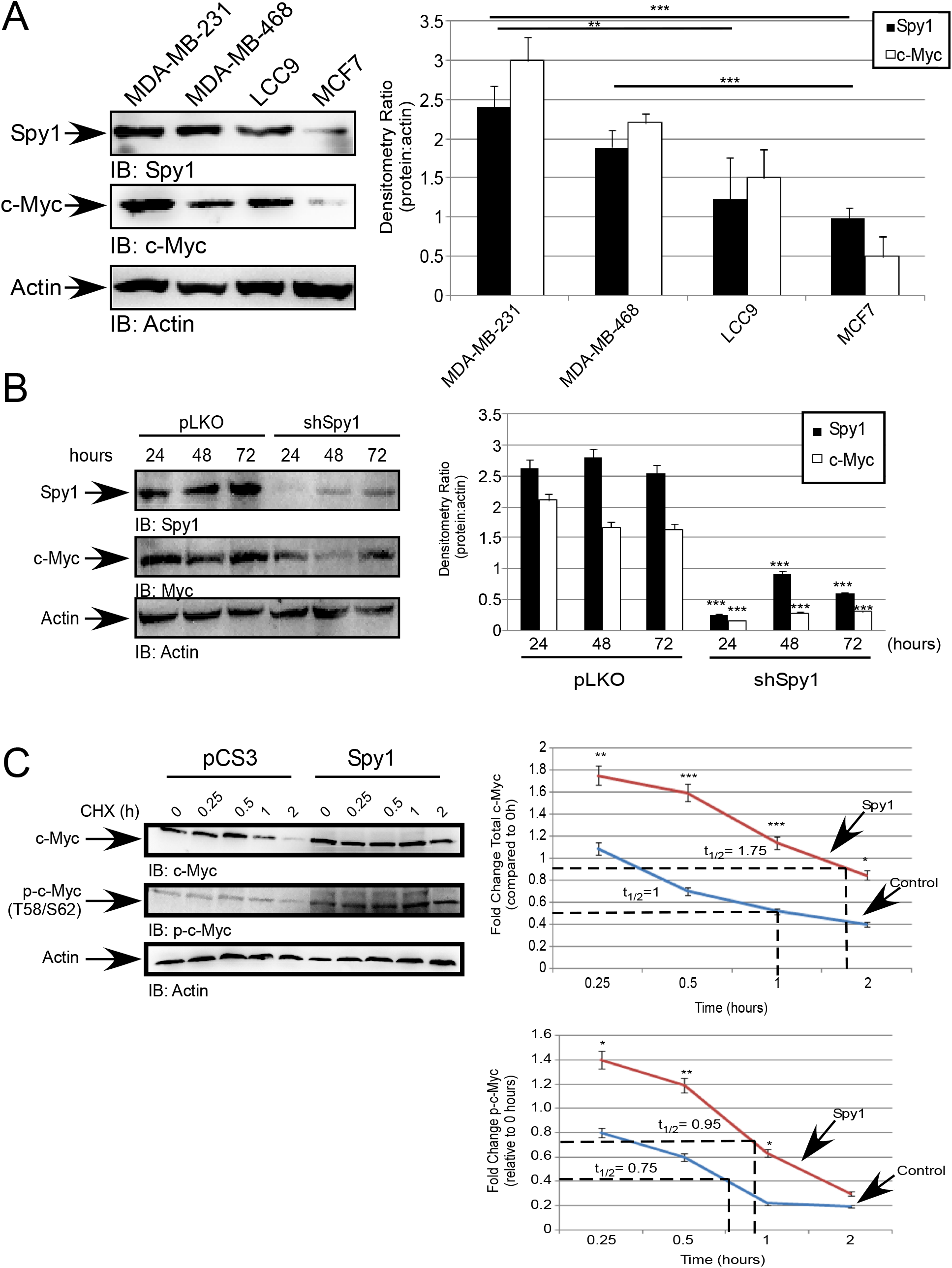
Spy1 has a role in stabilizing c-Myc. (A) A panel of breast cancer cell lines were subjected to protein extraction and SDS-PAGE analysis followed by IB. (B) MDA-MB-231 cells were infected with indicated constructs (along top of representative blot and x-axis of densitometry graph) (C) Hek-293 cells were transfected with indicated constructs (along top of representative blot and x-axis of densitometry graph); followed by treatment with 50 ug/ml cyclohexamide. After 0.25-2 hrs, cells were subjected to SDS-PAGE analysis and IB. Fold change of control and overexpression to time 0 hours was calculated and graphed to determine the overall effect on c-Myc stabilization by overexpression of Spy1. (A-C) Error bars reflect SE between triplicate experiments. Student’s t-test was performed; *p<0.05, **p<0.01,***p<0.001.

### Stabilization of c-Myc requires Spy1 and Cyclin E

c-Myc protein levels are, in part, regulated post-translationally via phosphorylation on serine (S)-62 leading to subsequent protein stabilization. S62 can be phosphorylated by CDK2, CDK1, and ERK1/2 [47-50]. To determine the effect of Spy1 on the stabilization of c-Myc, Spy1 was overexpressed in HEK-293 cells and Cyclin E1 was used as a positive control. Our data confirms the literature that Cyclin E/CDK2 results in an increased phosphorylation of c-Myc at S62 (Figure 4A). Comparably, we show that overexpression of Spy1 also leads to the phosphorylation of c-Myc (Figure 4A). We further investigated whether Spy1 was a necessary mediator of c-Myc stabilization by knocking down either Spy1 or Cyclin E1 in HEK-293 cells. Knockdown of either gene in HEK-293 significantly decreased the level of p-c-Myc as compared to control (Figure 4B).

**Figure 4.**
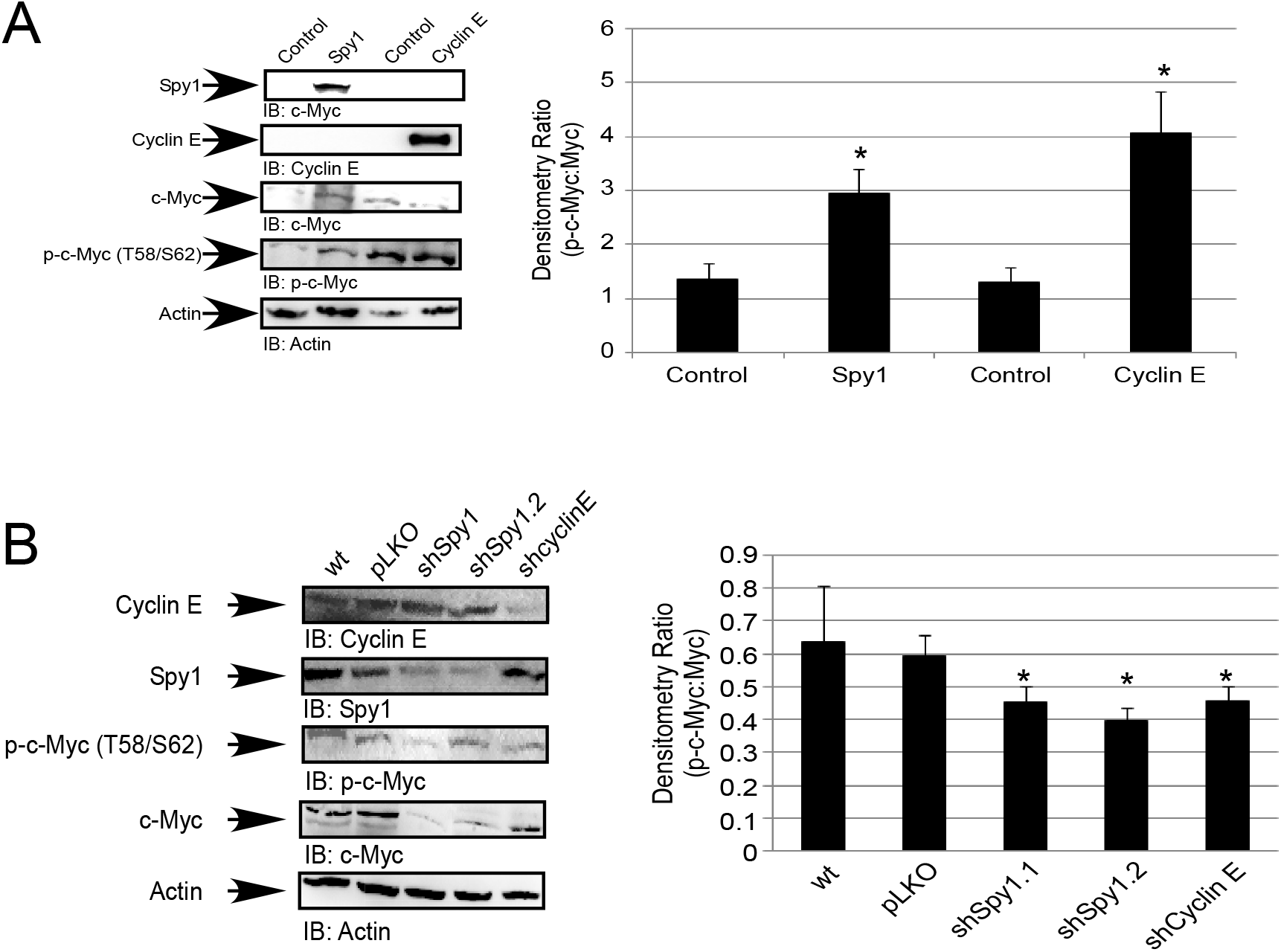
Stabilization of c-Myc requires Spy1 and Cyclin E. (A) Cells were transfected with indicated constructs; followed by SDS-PAGE and IB. (B) Cells were infected with indicated constructs; followed by SDS-PAGE and IB. (A-B) A representative blot is shown (left panel). Densitometry ratio of protein to actin loading control is shown (right panel). Error bars reflect SE between triplicate experiments. Student’s t-test was performed; *p<0.05.

### Spy1 levels are elevated in human TNBC tumour tissue

Spy1 levels are elevated in invasive ductal carcinoma of the breast [33]. To determine the levels of Spy1 in TNBC, frozen adjacent pair-matched normal and tumour TNBC tumour samples were obtained from the Ontario Tumour Bank and subjected to protein extraction and SDS-PAGE analysis. Spy1 levels are significantly elevated in TNBC patient samples as compared to adjacent pair-matched normal tissue (Figure 5A). To increase our sample size TNBC embedded patient samples were collected, subject to haematoxylin and eosin (H&E) staining to designate tumour versus stroma area and 1.5 mm cores used in a TMA construction. At least 3 normal patient samples were included on each array and IHC/DAB staining was performed for Spy1 (Figure 5B-C) and c-Myc protein levels (Figure 5D-E). Spy1 protein levels are significantly higher than in control, with an increase in intensity of over 2-fold over all samples (Figure 5C). c-Myc protein levels were also significantly elevated as compared to normal tissue (Figure 5E).

**Figure 5.**
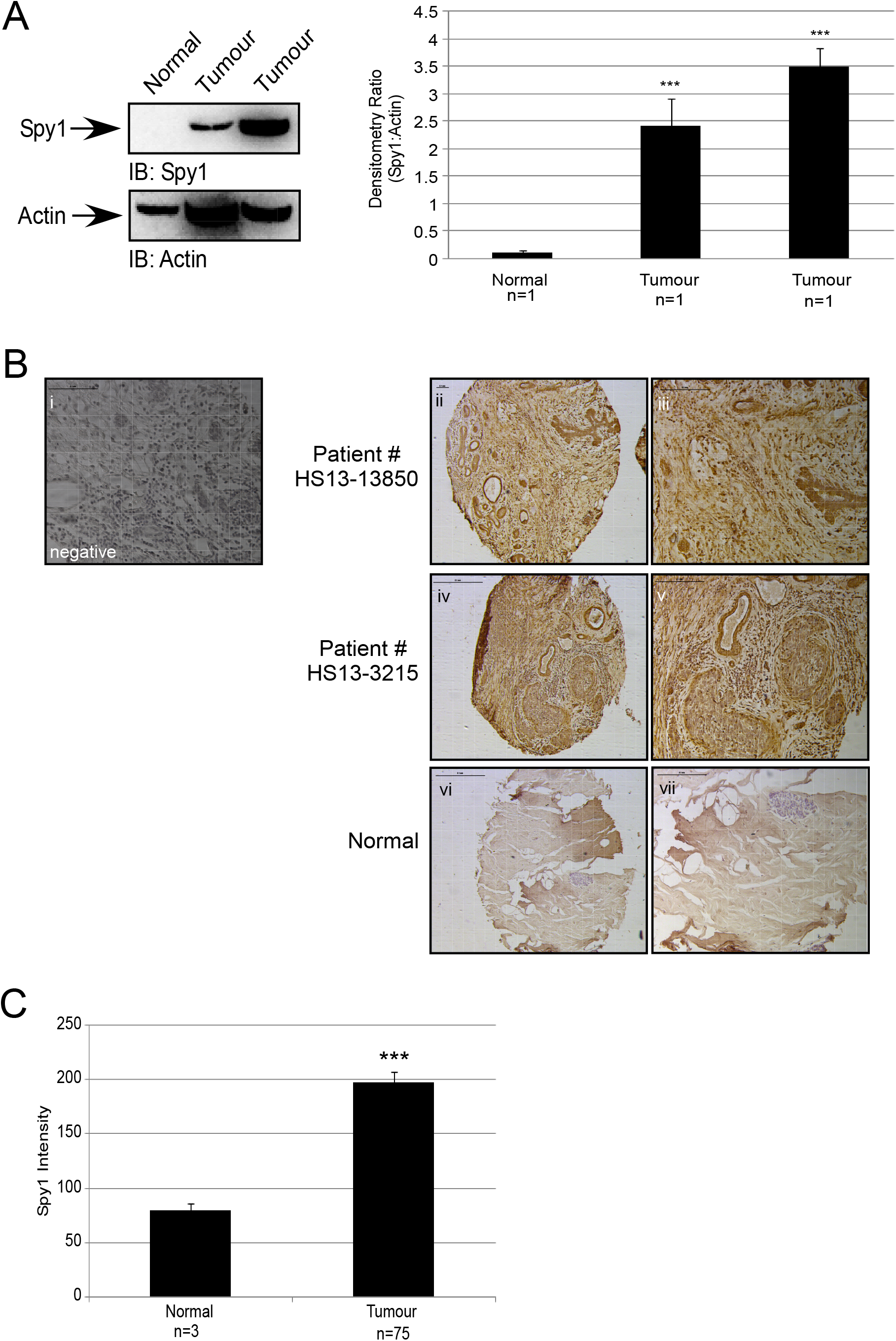
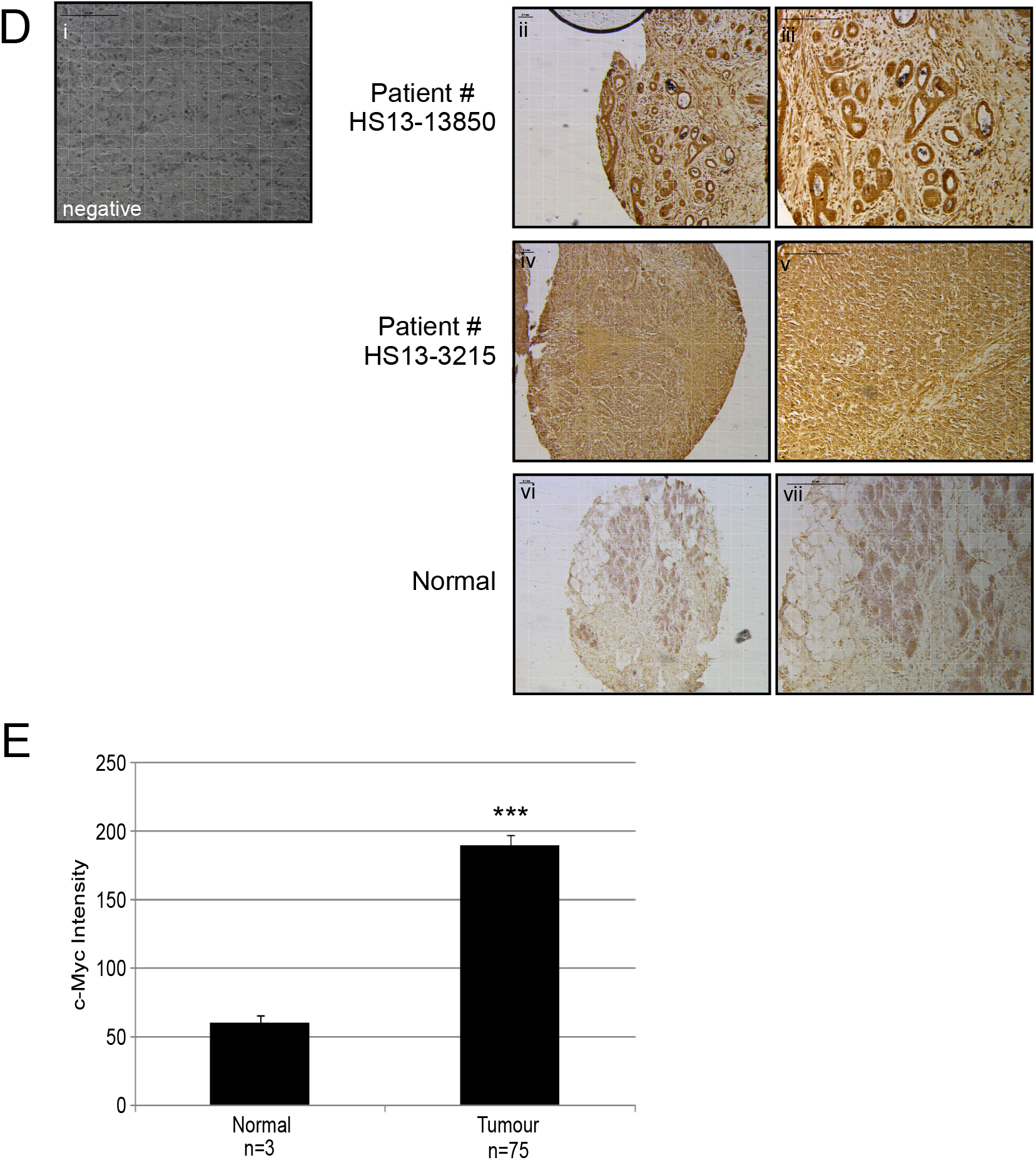
Spy1 levels elevated in human TNBC tumour tissue. (A) Protein from frozen tumour tissue was extracted and analysed by SDS-PAGE and IB. Densitometry analysis for Spy1 levels is seen in the right panel. TMAs were constructed and stained for Spy1 protein (B-C) or c-Myc protein (D-E). (B) Representative images of patient samples stained for Spy1 protein. (i) negative control. (ii, iv & vi) Image taken at 79.7x using a 1x stereoscope. (iii, v & vii) Image taken at 159x using a 2x stereoscope. (C) Average Spy1 intensity over 60 patient samples obtained by random sampling of 10×10 pixels each and quantified in Adobe Photoshop. (D) Representative images of samples stained for c-Myc protein. (i) negative control. (ii, iv & vi) Image taken at 79.7x using a 1x stereoscope. (iii, v & vii) Image taken at 159x using a 2x stereoscope. (E) Average c-Myc intensity over 60 patient samples obtained by random sampling of 10×10 pixels each and quantified in Adobe Photoshop. (A, C, & E) Error bars reflect SE between triplicate cores. Student’s t-test was performed; ***p<0.001.

### Spy1 knockdown increases TNBC cell line response to chemotherapy treatment

ERα-negative breast cancers, including TNBC, undergo a common chemotherapy regimen which includes an anthracycline, cyclophosphamide, and taxol combination, also known as AC/T [51]. To determine whether chemotherapy regimens can work more effectively *in vitro* when Spy1 levels have been depleted or are low, MDA-MB-231 cells were infected to knockdown either Spy1 or Cyclin E1 (Figure 6A). Knockdown of either gene significantly reduced levels of c-Myc. However, Spy1 knockdown also demonstrated a significant decrease in percent cell viability with the use of each drug treatment alone or in combination (AC/T). This effect was not consistently seen with Cyclin E1 knockdown, especially with the individual use of paclitaxel (Figure 6B), indicating this is a trait unique to Spy1.

**Figure 6.**
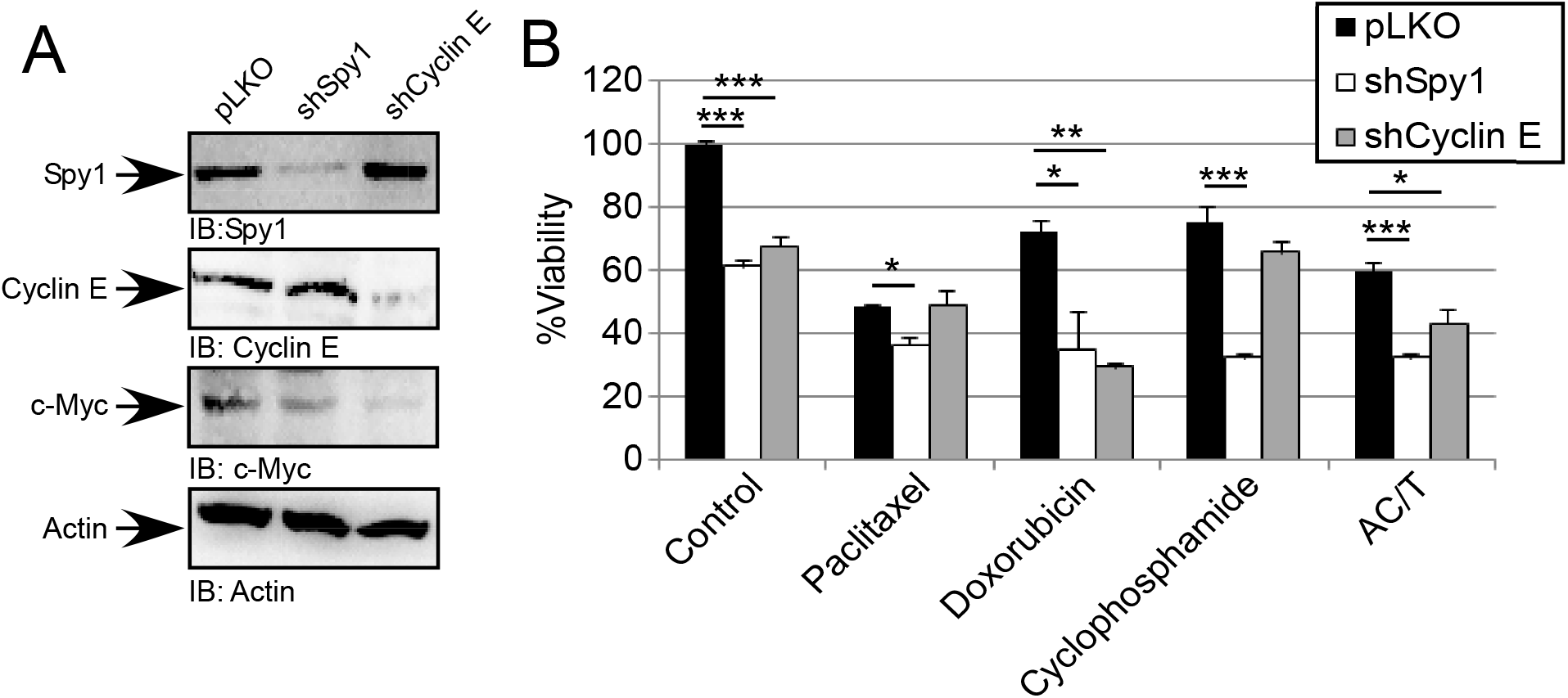
Spy1 knockdown increases TNBC cell line response to chemotherapy treatment. Cells were infected using indicated constructs. (A) Confirmation of knockdown was seen through SDS-PAGE and IB analysis. (B) Cells were treated with 100 nM paclitaxel, 25 nM doxorubicin, 6 mM cyclophosphamide, or a combination of the three (AC/T). Following incubation times, cells were subjected to trypan blue exclusion assay. Error bars reflect SE between triplicate experiments. Two-way ANOVA was performed; *p<0.05, **p<0.01, ***p<0.001.

## DISCUSSION

Breast cancer is a heterogenous disease, divided into 5 main subtypes. Each subtype is defined by its molecular signature; with 3 important proteins to aid in its classification; ERα, PR, and Her2/neu [52]. The presence of at least one of these three proteins enables a patient to receive targeted therapies; which have significantly increased the overall 5-year survival rate to 88% [53]. However, a subset of patients cannot respond to these forms of treatments or become resistant to therapies through the loss of ERα, PR, or Her2/neu [52]. Loss/absence of all three genes is classified as the highly aggressive TNBC group, with a majority of patients being under 50 years of age [52]. The aggressiveness of this subtype is due in part to the upregulation of various genes, including the tumour suppressor gene p53 and the oncogene, c-Myc [1, 21].

Transcriptional upregulation of c-Myc over time can confer control over ERα-targeted genes [21, 22]. This characteristic of c-Myc has been correlated with the basal breast cancer subtype and acquired resistance to hormone therapies, such as tamoxifen [54-56]. Our data shows that persistent c-Myc signalling using MMTV-Myc cells results in decreased response to tamoxifen over time, leading to a resistant phenotype; correlating with a loss of ERα protein expression. The mechanism behind this differential response and loss of expression is currently unknown. There is data to suggest that loss/downregulation of ERα may be through the activation of downstream c-Myc targets that feedback to ERα [54]. Much research has focused on targeting c-Myc; however, this has proven challenging due to the vast number of c-Myc targets, directing multiple biological functions [23, 24]. c-Myc null cells have a 2.5 fold decrease in protein synthesis which leads to growth defects due to a delay in both G1 and G2 phases of the cell cycle [20]. Whether targeting specific downstream targets, such as the atypical cyclin Spy1, can avoid lethality of healthy cells and provide adequate targeting to cancer cells is a hypothesis that this work supports.

Spy1 has been found to follow a similar expression profile as c-Myc within the developing mammary gland [38], and we showed Spy1 protein levels remain upregulated in c-Myc driven breast cancer cells in culture as they acquire resistance to hormone therapy. Spy1 and c-Myc are upregulated in similar cancers, such as neuroblastoma and invasive breast carcinomas [1, 2, 10, 13, 33, 34, 37, 38]. We show, for the first time, that Spy1 and c-Myc are co-regulated in TNBC patient samples and cell lines and that Spy1 levels influence the protein stabilization of c-Myc. Importantly, in c-Myc overexpressing MMTV cells, knockdown of Spy1 significantly decreases the rate of proliferation, and sensitized late passage resistant cells to tamoxifen. ERα-negative tumours, specifically TNBC, are known to have no response to hormone therapies and are dependent on a chemotherapeutic regimen [14, 57]. Knockdown of Spy1 in TNBC cells significantly sensitized cells to both single agent as well as combination AC/T therapy. Importantly, the sensitivity seen with Spy1 knockdown was unique; similar effects were not seen in all treatments when Cyclin E1 was knocked down. We have shown that although Spy1 functions similar to Cyclin E1 in stabilizing and influencing the levels of c-Myc, manipulation of Spy1 levels uniquely enhances sensitivity to treatment. Spy1 has a unique set of substrates and overrides CDKs using a mechanism different than that of classical cyclin-CDKs [30, 58, 59]; hence, targeting Spy1-driven CDKs in cancers with high expression levels of both c-Myc and Spy1 could promote novel, and potentially specific therapeutics.

## Conclusions

We have shown high levels of Spy1 alter the responsiveness of TNBC cells to chemotherapeutic and hormone therapies. Furthermore, we show, for the first time, that increased Spy1 protein levels correlate with high c-Myc protein levels in TNBC patient tumour samples. Importantly, targeting Spy1 in cancers with high levels of c-Myc can restore a response to therapeutics. In summary, we show Spy1 may act as a novel therapeutic target in the clinic.

## List of Abbreviations

BL: basal-like
BSA: bovine serum albumin
CDK: cyclin dependent kinase
CHX: cyclohexamide
ERα: estrogen receptor alpha
Her2/neu: human epidermal growth factor receptor 2
IB: immunoblotting
IM: immunomodulatory
IHC: immunohistochemistry
LAR: luminal androgen receptor
M: mesenchymal
MMTV: mouse mammary tumour virus
MSL: mesenchymal stem-like
PBS: phosphate buffered saline
PR: progesterone receptor
Spy1: Speedy
TMA: tissue microarray
TNBC: triple negative breast cancer

## Authors’ Contributions

R-MF carried out all of the experiments described herein and aided in the draft of the manuscript. BF aided in the draft of the manuscript. CH is a valuable collaborator and the principal investigator in the clinic, acquiring patient consent and tissue for the trial. LAP funded the project and had a lead role in study design, interpretation of the data and manuscript preparation. All authors edited the manuscript and provided comments on the intellectual content.

## Acknowledgments

We would like to thank the Windsor Regional Cancer Centre, pathology department, and Dr. D. Shum for the assistance in acquiring patient tumour samples. We acknowledge and thank the patients of the clinical trial. Biological Materials were provided by the Ontario Tumour Bank, which is funded by the Ontario Institute for Cancer Research.

## Ethics Approval

Approval to receive samples from the Ontario Tumour Bank and Windsor Regional Hospital has been approved by the University of Windsor Research Ethics Board under REB #14-123.

## REFERENCES

1. Perou CM: Molecular stratification of triple-negative breast cancers. Oncologist 2010, 15 Suppl 5:39–48.

2. Perou CM, Sorlie T, Eisen MB, van de Rijn M, Jeffrey SS, Rees CA, Pollack JR, Ross DT, Johnsen H, Akslen LA et al: Molecular portraits of human breast tumours. Nature 2000, 406(6797):747–752.

3. Prat A, Parker JS, Karginova O, Fan C, Livasy C, Herschkowitz JI, He X, Perou CM: Phenotypic and molecular characterization of the claudin-low intrinsic subtype of breast cancer. Breast Cancer Res 2010, 12(5):R68.

4. Malhotra GK, Zhao X, Band H, Band V: Histological, molecular and functional subtypes of breast cancers. Cancer Biol Ther 2010, 10(10):955–960.

5. Lehmann BD, Bauer JA, Chen X, Sanders ME, Chakravarthy AB, Shyr Y, Pietenpol JA: Identification of human triple-negative breast cancer subtypes and preclinical models for selection of targeted therapies. J Clin Invest 2011, 121(7):2750–2767.

6. Lumachi F, Brunello A, Maruzzo M, Basso U, Basso SM: Treatment of estrogen receptor-positive breast cancer. Curr Med Chem 2013, 20(5):596–604.

7. Lindström LS, Karlsson E, Wilking UM, Johansson U, Hartman J, Lidbrink EK, Hatschek T, Skoog L, Bergh J: Clinically Used Breast Cancer Markers Such As Estrogen Receptor, Progesterone Receptor, and Human Epidermal Growth Factor Receptor 2 Are Unstable Throughout Tumor Progression. Journal of Clinical Oncology 2012, 30(21):2601–2608.

8. Yang X, Phillips DL, Ferguson AT, Nelson WG, Herman JG, Davidson NE: Synergistic activation of functional estrogen receptor (ER)-alpha by DNA methyltransferase and histone deacetylase inhibition in human ER-alpha-negative breast cancer cells. Cancer Res 2001, 61(19):7025–7029.

9. Escot C, Theillet C, Lidereau R, Spyratos F, Champeme MH, Gest J, Callahan R: Genetic alteration of the c-myc protooncogene (MYC) in human primary breast carcinomas. Proc Natl Acad Sci U S A 1986, 83(13):4834–4838.

10. Kniazev PG, Schafer R, Willecke K, Pluzhnikova GF, Serova OM: [Activation of ras and myc proto-oncogenes in human breast carcinoma and neuroblastoma]. Mol Biol (Mosk) 1986, 20(5):1236–1243.

11. Kozbor D, Croce CM: Amplification of the c-myc oncogene in one of five human breast carcinoma cell lines. Cancer Res 1984, 44(2):438–441.

12. Persons DL, Borelli KA, Hsu PH: Quantitation of HER-2/neu and c-myc gene amplification in breast carcinoma using fluorescence in situ hybridization. Mod Pathol 1997, 10(7):720–727.

13. Xu J, Chen Y, Olopade OI: MYC and Breast Cancer. Genes Cancer 2010, 1(6):629–640.

14. Peddi PF, Ellis MJ, Ma C: Molecular basis of triple negative breast cancer and implications for therapy. Int J Breast Cancer 2012, 2012:217185.

15. Kang J, Sergio CM, Sutherland RL, Musgrove EA: Targeting cyclin-dependent kinase 1 (CDK1) but not CDK4/6 or CDK2 is selectively lethal to MYC-dependent human breast cancer cells. BMC Cancer 2014, 14:32.

16. Bieche I, Laurendeau I, Tozlu S, Olivi M, Vidaud D, Lidereau R, Vidaud M: Quantitation of MYC gene expression in sporadic breast tumors with a real-time reverse transcription-PCR assay. Cancer Res 1999, 59(12):2759–2765.

17. Blakely CM, Sintasath L, D’Cruz CM, Hahn KT, Dugan KD, Belka GK, Chodosh LA: Developmental stage determines the effects of MYC in the mammary epithelium. Development 2005, 132(5):1147–1160.

18. Liao DJ, Dickson RB: c-Myc in breast cancer. Endocr Relat Cancer 2000, 7(3):143–164.

19. Littlewood TD, Evan GI: The role of myc oncogenes in cell growth and differentiation. Adv Dent Res 1990, 4:69–79.

20. Schmidt EV: The role of c-myc in cellular growth control. Oncogene 1999, 18(19):2988–2996.

21. Alles MC, Gardiner-Garden M, Nott DJ, Wang Y, Foekens JA, Sutherland RL, Musgrove EA, Ormandy CJ: Meta-analysis and gene set enrichment relative to er status reveal elevated activity of MYC and E2F in the “basal” breast cancer subgroup. PLoS ONE 2009, 4(3):e4710.

22. Dadiani M, Seger D, Kreizman T, Badikhi D, Margalit R, Eilam R, Degani H: Estrogen regulation of vascular endothelial growth factor in breast cancer in vitro and in vivo: the role of estrogen receptor alpha and c-Myc. Endocr Relat Cancer 2009, 16(3):819–834.

23. Amati B, Land H: Myc-Max-Mad: a transcription factor network controlling cell cycle progression, differentiation and death. Curr Opin Genet Dev 1994, 4(1):102–108.

24. Amati B, Littlewood TD, Evan GI, Land H: The c-Myc protein induces cell cycle progression and apoptosis through dimerization with Max. EMBO J 1993, 12(13):5083–5087.

25. Evan G, Harrington E, Fanidi A, Land H, Amati B, Bennett M: Integrated control of cell proliferation and cell death by the c-myc oncogene. Philos Trans R Soc Lond B Biol Sci 1994, 345(1313):269–275.

26. Evan GI, Wyllie AH, Gilbert CS, Littlewood TD, Land H, Brooks M, Waters CM, Penn LZ, Hancock DC: Induction of apoptosis in fibroblasts by c-myc protein. Cell 1992, 69(1):119–128.

27. Soucek L, Jucker R, Panacchia L, Ricordy R, Tato F, Nasi S: Omomyc, a potential Myc dominant negative, enhances Myc-induced apoptosis. Cancer Res 2002, 62(12):3507–3510.

28. Horiuchi D, Kusdra L, Huskey NE, Chandriani S, Lenburg ME, Gonzalez-Angulo AM, Creasman KJ, Bazarov AV, Smyth JW, Davis SE et al: MYC pathway activation in triple-negative breast cancer is synthetic lethal with CDK inhibition. J Exp Med 2012, 209(4):679–696.

29. Nebreda AR: CDK activation by non-cyclin proteins. Curr Opin Cell Biol 2006, 18(2):192–198.

30. Cheng A, Gerry S, Kaldis P, Solomon MJ: Biochemical characterization of Cdk2-Speedy/Ringo A2 BMC Biochem 2005, 6:19.

31. McAndrew CW, Gastwirt RF, Meyer AN, Porter LA, Donoghue DJ: Spy1 enhances phosphorylation and degradation of the cell cycle inhibitor p27. Cell Cycle 2007, 6(15):1937–1945.

32. McGrath DA, Fifield BA, Marceau AH, Tripathi S, Porter LA, Rubin SM: Structural basis of divergent cyclin-dependent kinase activation by Spy1/RINGO proteins. EMBO J 2017, 36(15):2251–2262.

33. Al Sorkhy M, Ferraiuolo RM, Jalili E, Malysa A, Fratiloiu AR, Sloane BF, Porter LA: The cyclin-like protein Spy1/RINGO promotes mammary transformation and is elevated in human breast cancer. BMC Cancer 2012, 12:45.

34. Zucchi I, Mento E, Kuznetsov VA, Scotti M, Valsecchi V, Simionati B, Vicinanza E, Valle G, Pilotti S, Reinbold R et al: Gene expression profiles of epithelial cells microscopically isolated from a breast-invasive ductal carcinoma and a nodal metastasis. Proc Natl Acad Sci U S A 2004, 101(52):18147–18152.

35. Hang Q, Fei M, Hou S, Ni Q, Lu C, Zhang G, Gong P, Guan C, Huang X, He S: Expression of Spy1 protein in human non-Hodgkin’s lymphomas is correlated with phosphorylation of p27 Kip1 on Thr187 and cell proliferation. Med Oncol 2012, 29(5):3504–3514.

36. Ke Q, Ji J, Cheng C, Zhang Y, Lu M, Wang Y, Zhang L, Li P, Cui X, Chen L et al: Expression and prognostic role of Spy1 as a novel cell cycle protein in hepatocellular carcinoma. Exp Mol Pathol 2009, 87(3):167–172.

37. Lubanska D, Porter LA: The atypical cell cycle regulator Spy1 suppresses differentiation of the neuroblastoma stem cell population. Oncoscience 2014, 1(5):336–348.

38. Golipour A, Myers D, Seagroves T, Murphy D, Evan GI, Donoghue DJ, Moorehead RA, Porter LA: The Spy1/RINGO family represents a novel mechanism regulating mammary growth and tumorigenesis. Cancer Res 2008, 68(10):3591–3600.

39. Huang Y, Liu Y, Chen Y, Yu X, Yang J, Lu M, Lu Q, Ke Q, Shen A, Yan M: Peripheral nerve lesion induces an up-regulation of Spy1 in rat spinal cord. Cell Mol Neurobiol 2009, 29(3):403–411.

40. Porter LA, Dellinger RW, Tynan JA, Barnes EA, Kong M, Lenormand JL, Donoghue DJ: Human Speedy: a novel cell cycle regulator that enhances proliferation through activation of Cdk2. J Cell Biol 2002, 157(3):357–366.

41. Barnes EA, Porter LA, Lenormand JL, Dellinger RW, Donoghue DJ: Human Spy1 promotes survival of mammalian cells following DNA damage. Cancer Res 2003, 63(13):3701–3707.

42. Gastwirt RF, Slavin DA, McAndrew CW, Donoghue DJ: Spy1 expression prevents normal cellular responses to DNA damage: inhibition of apoptosis and checkpoint activation. J Biol Chem 2006, 281(46):35425–35435.

43. Hamm C FB, Kay A, Kulkarni S, Gupta R, Mathews J, Ferraiuolo RM, Al-Wahsh H, Mailloux E, Hussein A, Porter LA: A prospective phase II clinical trial identifying the optimal regimen for carboplatin plus standard backbone of anthracycline and taxane-based chemotherapy in triple negative breast cancer. Medical Oncology 2022, 39(49).

44. Goenka S, Peelukhana SV, Kim J, Stringer KF, Banerjee RK: Dependence of vascular damage on higher frequency components in the rat-tail model. Ind Health 2013, 51(4):373–385.

45. Matkowskyj KA, Schonfeld D, Benya RV: Quantitative immunohistochemistry by measuring cumulative signal strength using commercially available software photoshop and matlab. J Histochem Cytochem 2000, 48(2):303–312.

46. Hydbring P, Bahram F, Su Y, Tronnersjo S, Hogstrand K, von der Lehr N, Sharifi HR, Lilischkis R, Hein N, Wu S et al: Phosphorylation by Cdk2 is required for Myc to repress Ras-induced senescence in cotransformation. Proc Natl Acad Sci U S A 2010, 107(1):58–63.

47. Adhikary S, Eilers M: Transcriptional regulation and transformation by Myc proteins. Nat Rev Mol Cell Biol 2005, 6(8):635–645.

48. Amati B: Myc degradation: dancing with ubiquitin ligases. Proc Natl Acad Sci U S A 2004, 101(24):8843–8844.

49. Amati B, Alevizopoulos K, Vlach J: Myc and the cell cycle. Front Biosci 1998, 3:d250–268.

50. Sears R, Nuckolls F, Haura E, Taya Y, Tamai K, Nevins JR: Multiple Rasdependent phosphorylation pathways regulate Myc protein stability. Genes Dev 2000, 14(19):2501–2514.

51. Citron ML, Berry DA, Cirrincione C, Hudis C, Winer EP, Gradishar WJ, Davidson NE, Martino S, Livingston R, Ingle JN et al: Randomized trial of dose-dense versus conventionally scheduled and sequential versus concurrent combination chemotherapy as postoperative adjuvant treatment of node-positive primary breast cancer: first report of Intergroup Trial C9741/Cancer and Leukemia Group B Trial 9741. J Clin Oncol 2003, 21(8):1431–1439.

52. Dolle JM, Daling JR, White E, Brinton LA, Doody DR, Porter PL, Malone KE: Risk factors for triple-negative breast cancer in women under the age of 45 years. Cancer Epidemiol Biomarkers Prev 2009, 18(4):1157–1166.

53. Siegel R, DeSantis C, Virgo K, Stein K, Mariotto A, Smith T, Cooper D, Gansler T, Lerro C, Fedewa S et al: Cancer treatment and survivorship statistics, 2012. CA Cancer J Clin 2012, 62(4):220–241.

54. Dimitrakakis C, Zhou J, Wang J, Matyakhina L, Mezey E, Wood JX, Wang D, Bondy C: Co-expression of estrogen receptor-alpha and targets of estrogen receptor action in proliferating monkey mammary epithelial cells. Breast Cancer Res 2006, 8(1):R10.

55. Musgrove EA, Sergio CM, Anderson LR, Inman CK, McNeil CM, Alles MC, Gardiner-Garden M, Ormandy CJ, Butt AJ, Sutherland RL: Identification of downstream targets of estrogen and c-myc in breast cancer cells. Adv Exp Med Biol 2008, 617:445–451.

56. Musgrove EA, Sergio CM, Loi S, Inman CK, Anderson LR, Alles MC, Pinese M, Caldon CE, Schutte J, Gardiner-Garden M et al: Identification of functional networks of estrogen-and c-Myc-responsive genes and their relationship to response to tamoxifen therapy in breast cancer. PLoS One 2008, 3(8):e2987.

57. Dent R, Trudeau M, Pritchard KI, Hanna WM, Kahn HK, Sawka CA, Lickley LA, Rawlinson E, Sun P, Narod SA: Triple-negative breast cancer: clinical features and patterns of recurrence. Clin Cancer Res 2007, 13(15 Pt 1):4429–4434.

58. Gastwirt RF, McAndrew CW, Donoghue DJ: Speedy/RINGO regulation of CDKs in cell cycle, checkpoint activation and apoptosis. Cell Cycle 2007, 6(10):1188–1193.

59. Karaiskou A, Perez LH, Ferby I, Ozon R, Jessus C, Nebreda AR: Differential regulation of Cdc2 and Cdk2 by RINGO and cyclins J Biol Chem 2001, 276(38):36028–36034.

